# AgamPrimer: Primer Design in *Anopheles gambiae* informed by range-wide genomic variation

**DOI:** 10.1101/2022.12.31.521737

**Authors:** Sanjay Curtis Nagi, Alistair Miles, Martin J Donnelly

## Abstract

The major malaria mosquito, *Anopheles gambiae s.l*, is one of the most studied organisms in medical research and also one of the most genetically diverse. When designing polymerase chain reaction (PCR) or hybridisation-based molecular assays, reliable primer and probe design is crucial. However, single nucleotide polymorphisms (SNPs) in primer binding sites can prevent primer binding, leading to null alleles, or bind suboptimally, leading to preferential amplification of specific alleles. Given the extreme genetic diversity of *An. gambiae*, researchers need to consider this genetic variation when designing primers and probes to avoid amplification problems. In this note, we present a python package, AgamPrimer, which exploits the AG1000G dataset and allows users to rapidly design primers in *An. gambiae*, whilst summarising genetic variation in the primer binding sites and visualising the position of primer pairs. AgamPrimer allows the design of both genomic DNA and cDNA primers and hybridisation probes. By coupling this python package with Google Colaboratory, AgamPrimer is an open and accessible platform for primer and probe design, hosted in the cloud for free. AgamPrimer is available here https://github.com/sanjaynagi/AgamPrimer and we hope it will be a useful resource for the community to design probe and primer sets that can be reliably deployed across the *An. gambiae* species range.

## Introduction

The polymerase chain reaction (PCR) is ubiquitous in molecular biology, providing template sequence for a wide array of techniques, such as detecting the presence or absence of particular DNA sequences, quantifying the abundance of transcripts, or in Sanger and next-generation sequencing. Primers - short, single-strand DNA sequences which bind to the template and facilitate amplification - are crucial to effective PCR reactions and must be designed to be robust, reliable and consistent across experimental conditions.

Single nucleotide polymorphisms (SNPs) in primer binding sites can affect both the stability of the primer-template duplex, as well as the efficiency with which DNA polymerases can extend the primer (Letowski, Brousseau, and Masson 2004; Wu, Hong, and Liu 2009). In some cases, this can completely prevent primer binding and amplification of the template DNA, often referred to as null alleles or allelic dropout (Carlson et al. 2006). On most genotyping platforms, these alleles are problematic and difficult to detect, as null allele heterozygotes will be indistinguishable from true homozygous individuals. Allelic dropout is known to cause problems in human genetic testing (Silva et al. 2017; Zajícková, Krepelová, and Zofková 2003). Null allele homozygotes could be suggested if a sample repeatedly fails to amplify, however, when performing PCR on pooled samples we would not observe this failure, and therefore can never know whether all samples amplified successfully. Ensuring genetic markers do not violate Hardy-Weinberg equilibrium (HWE) is one way to partially safeguard against this problem (Chapuis and Estoup 2007), however, this is not always performed in practice, and excluding such markers may lead to loss of information when HWE deviation has another cause.

Another problematic scenario occurs if primers do bind but with unequal efficiency against different genetic variants. In this case, any molecular assay that is quantitative, such as qPCR for gene expression, could be severely affected and lead to biases in the estimation of sequence abundance between genetic variants or strains (Lefever et al. 2013). A previous study found that single mismatches can introduce a range of impacts on Cq values, ranging from relatively minor (<1.5) to major (>7.0) (Stadhouders et al. 2010). The impact of a variant on primer binding depends on multiple factors but mismatches within the last 5 nucleotides at the 3’ end can disrupt the nearby polymerase active site, and so these mismatches tend to have a much greater impact (Stadhouders et al. 2010; Martins et al. 2011). Primers should therefore be designed to avoid these sites or if unavoidable, to contain degenerate bases at the sites of SNPs, in order to maximise the robustness of molecular experiments (Quinlan and Marth 2007).

The *Anopheles gambiae* 1000 genomes project has revealed staggering amounts of genetic variation in the major malaria mosquito, *Anopheles gambiae s.l (Miles et al. 2017)* with a segregating SNP in less than every 2 bases of the accessible genome (The Anopheles gambiae 1000 Genomes Consortium 2020). Despite this, the vast majority of primers designed to target the *Anopheles gambiae s.l* genome do not consider SNP variation. In the past, this was not straightforward, as it would require both handling large genomic datasets and matching designed primers to genomic positions. Thanks to recent advances in cloud computing and the malariagen_data API, we can now design primers in the cloud whilst checking for genetic variation in the *Anopheles gambiae* 1000 genomes project. In this note we present AgamPrimer, a python package which is coupled with a Google Colaboratory notebook, allowing users to easily design primers and probes in the cloud whilst considering genetic variation in *Anopheles gambiae s.l*.

## Methods and Implementation

AgamPrimer is a two-phase process, first designing sets of primers and probes and secondly investigating SNP variation in the targeted sites. AgamPrimer uses Primer3 as the core primer design engine, in the form of Primer3-py. Primer3 is open-source and has become the *de-facto* standard for primer design for molecular biology. Primer3-py is a set of recently developed python bindings for the Primer3 program (Untergasser et al. 2012), which can be run readily in a Google Colaboratory environment. To load genetic variation data from the *Anopheles* 1000 genomes project, we integrate the malariagen_data API, which allows rapid download and analysis of genomic data from the cloud. Integration of the PyData stack in malariagen_data allows users to perform rapid genomic analysis on large datasets where compute resources are modest, such as in Google Colaboratory notebooks. Google Colaboratory is a proprietary version of Jupyter Notebook and is provided for free alongside CPU and GPU access to anyone with a Google account.

AgamPrimer can be run in two ways, either running the full Colaboratory notebook in a step-wise fashion, or in a single command which produces all outputs, which may be preferred in more high-throughput primer design settings. Users may select primer design parameters by providing a python dictionary, or the primer3 default parameters can be used.

### Primer Design with *primer3*

AgamPrimer allows the design of genomic DNA primers, hybridisation probes or cDNA primers (for gene expression purposes). In the case of cDNA primers, one of the forward or reverse primers will be designed to span an exon-exon junction where available, to prevent the amplification of genomic DNA in the sample.

Table 1 shows the output from the initial phase of primer design. AgamPrimer reports the primer sequences, along with information on melting point, GC content, amplicon size and position in the target sequence, though the full Primer3 output is accessible to the user. The user may specify the number of desired primer pairs to design. After the Primer3 run, AgamPrimer will print out run statistics which may be useful for troubleshooting.

**Table 1.**
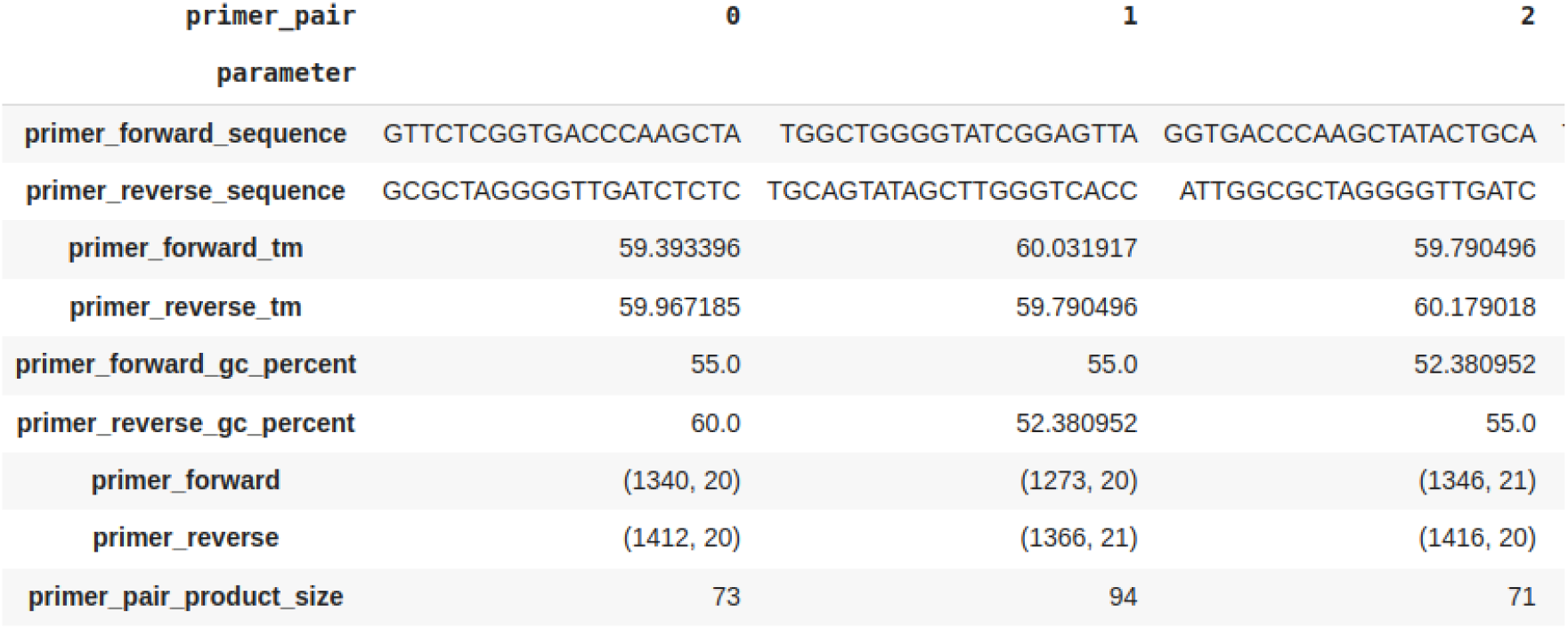
Primer3 results: A pandas dataframe generated by AgamPrimer. Useful information from each primer set is stored, such as the sequence, melting temperature and GC content.

### Interrogating the ag3 resource

The malariagen_data python package pulls in Ag1000g data from the cloud, facilitating rapid analysis of over 10,000 *An. gambiae s.l* whole genomes from throughout sub-Saharan Africa. In step 1 of the primer design process, we record the genomic positions of the designed primers, and in step 2 use these coordinates to extract SNP allele frequency information for given Ag1000g samples of choice. In the Colaboratory notebook, we generate a summary table of the Ag1000g inventory, counting samples by taxon, sample set and country, to guide users in selecting an appropriate cohort. Through the use of sample set identifiers, and sample queries (following standard pandas syntax), users may select any group of samples in the dataset to interrogate. Alternatively, the default settings will use every available mosquito genome. A sample query can be performed on any column of the sample metadata, such as species (taxon), country, year or location, amongst other metadata.

We then generate an interactive plot (Figure 1) which shows SNP variation in designed primer binding sites, in the user-selected Ag1000g cohort. The user can hover over points, which returns the exact frequencies of each nucleotide at that genomic position, which may be useful in the case where the user would prefer to design degenerate primers, as opposed to avoiding that primer set entirely. The plot also highlights the 3’ and 5’ prime ends, as well as the genomic span, GC content and melting temperature, allowing the user to easily and rapidly identify suitable oligonucleotides.

**Figure 1.**
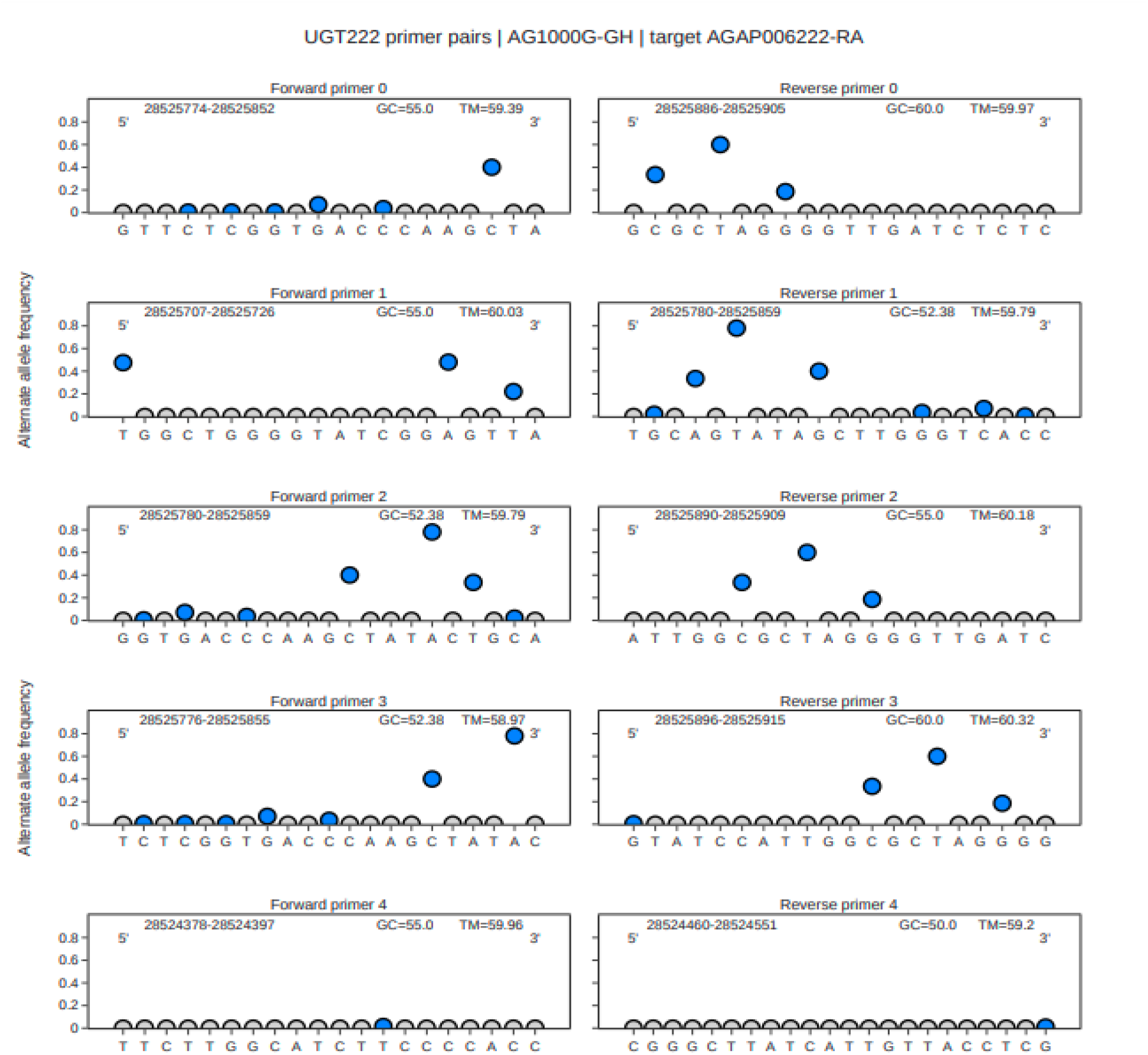
Illustrative plots showing allele frequencies in primer binding sites targeting the AGAP006222-RA transcript in specimens of *An. gambiae ss*. and *Anopheles coluzzii* from Ghana. An interactive plot with Plotly displays the primer or probe sequences from 5’ to 3’, with circles indicating the summed alternate allele frequency at that genomic position. Blue circles indicate segregating SNPs, and grey circles indicate sites which are invariant in the ag3 cohort of choice. The genomic span of each oligo is displayed alongside the GC content and Tm.

### Genomic location of primers

AgamPrimer then plots the position of the primer in the genome, in relation to any nearby exons. In Figure 2, we can see that all but one primer pair were designed at the Exon 4 and 5 boundary.

**Figure 2.**
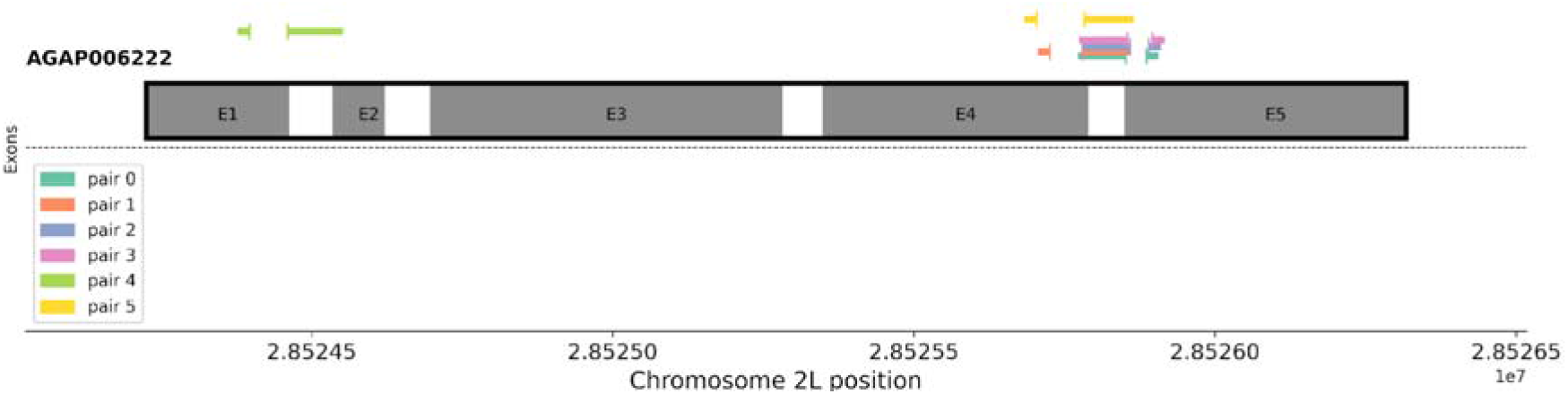
The genomic locations of designed primer sets in relation to nearby exons. Primers spanning exons are shown as expanded to clearly illustrate span of the whole junction for visualisation purposes, and only contain sequence at each extremity.

Primer pair 4, which targets the Exon 1 and 2 junction, contains much less SNP variation than the other primers.

To ensure the specificity of the designed primer and probes for only one genomic location, we align oligonucleotides to the AgamP3 genome with BLAT, using the gget python package API.

## Discussion

AgamPrimer integrates Primer3-py and the malariagen_data API to rapidly and conveniently design variation-informed primers and probes for molecular biology. Through the use of forms in Colaboratory, users are able to define their own parameters, which means that the AgamPrimer notebook does not require programming skills. This is an extremely important point, as we hope the tool will be useful for all researchers including molecular biologists who may not have programming experience.

Genomic surveillance of malaria mosquitoes is becoming increasingly important, with a number of high throughput amplicon sequencing panels having been developed to identify species across the entire *Anopheles* genus (Boddé et al. 2022; Makunin et al. 2022), within the *Anopheles gambiae* complex (Caputo et al. 2021), and to karyotype samples (Love et al. 2020). In the near future, it is likely that other amplicon sequencing panels will be designed to target phenotypes of interest, such as insecticide resistance, gene drive resistance, or vector competence. Just as in more standard genotyping assays, in amplicon sequencing, robust primer design is crucial, and therefore having a computational framework to design primers will prove invaluable.

AgamPrimer is open-source and free to use, and we encourage suggestions and contributions to the software. In the future, we plan to incorporate an API for another major malaria vector, *Anopheles funestus*, to further increase the utility of the package. The package is available here https://github.com/sanjaynagi/AgamPrimer and on the python package index (PyPi).

## References

Boddé, Marilou, Alex Makunin, Diego Ayala, Lemonde Bouafa, Abdoulaye Diabaté, Uwem Friday Ekpo, Mahamadi Kientega, et al. 2022. “High Resolution Species Assignment of Anopheles Mosquitoes Using K-Mer Distances on Targeted Sequences.” bioRxiv. https://doi.org/10.1101/2022.03.18.484650.

Caputo, Beniamino, Verena Pichler, Giordano Bottà, Carlo De Marco, Christina Hubbart, Eleonora Perugini, Joao Pinto, Kirk A. Rockett, Alistair Miles, and Alessandra Della Torre. 2021. “Novel Genotyping Approaches to Easily Detect Genomic Admixture between the Major Afrotropical Malaria Vector Species, Anopheles Coluzzii and An. Gambiae.” Molecular Ecology Resources 21 (5): 1504–16.

Carlson, Christopher S., Joshua D. Smith, Ian B. Stanaway, Mark J. Rieder, and Deborah A. Nickerson. 2006. “Direct Detection of Null Alleles in SNP Genotyping Data.” Human Molecular Genetics 15 (12): 1931–37.

Chapuis, Marie-Pierre, and Arnaud Estoup. 2007. “Microsatellite Null Alleles and Estimation of Population Differentiation.” Molecular Biology and Evolution 24 (3): 621–31.

Lefever, Steve, Filip Pattyn, Jan Hellemans, and Jo Vandesompele. 2013. “Single-Nucleotide Polymorphisms and Other Mismatches Reduce Performance of Quantitative PCR Assays.” Clinical Chemistry 59 (10): 1470–80.

Letowski, Jaroslaw, Roland Brousseau, and Luke Masson. 2004. “Designing Better Probes: Effect of Probe Size, Mismatch Position and Number on Hybridization in DNA Oligonucleotide Microarrays.” Journal of Microbiological Methods 57 (2): 269–78.

Love, R. Rebecca, Marco Pombi, Moussa W. Guelbeogo, Nathan R. Campbell, Melissa T. Stephens, Roch K. Dabire, Carlo Costantini, Alessandra Della Torre, and Nora J. Besansky. 2020. “Inversion Genotyping in the Anopheles Gambiae Complex Using High-Throughput Array and Sequencing Platforms.” G3 10 (9): 3299–3307.

Makunin, Alex, Petra Korleviċ, Naomi Park, Scott Goodwin, Robert M. Waterhouse, Katharina von Wyschetzki, Christopher G. Jacob, et al. 2022. “A Targeted Amplicon Sequencing Panel to Simultaneously Identify Mosquito Species and Plasmodium Presence across the Entire Anopheles Genus.” Molecular Ecology Resources 22 (1): 28–44.

Martins, Estefânia M., Laura Vilarinho, Sofia Esteves, Mónica Lopes-Marques, António Amorim, and Luísa Azevedo. 2011. “Consequences of Primer Binding-Sites Polymorphisms on Genotyping Practice.” Open Journal of Genetics 01 (02): 15–17.

Miles, Alistair, Nicholas J. Harding, Giordano Bottà, Chris S. Clarkson, Tiago Antão, Krzysztof Kozak, Daniel R. Schrider, et al. 2017. “Genetic Diversity of the African Malaria Vector Anopheles Gambiae.” Nature 552: 96–100.

Quinlan, Aaron R., and Gabor T. Marth. 2007. “Primer-Site SNPs Mask Mutations.” Nature Methods 4 (3): 192.

Silva, Felipe Carneiro, Giovana Tardin Torrezan, Rafael Canfield Brianese, Raquel Stabellini, and Dirce Maria Carraro. 2017. “Pitfalls in Genetic Testing: A Case of a SNP in Primer-Annealing Region Leading to Allele Dropout in BRCA1.” Molecular Genetics & Genomic Medicine 5 (4): 443–47.

Stadhouders, Ralph, Suzan D. Pas, Jeer Anber, Jolanda Voermans, Ted H. M. Mes, and Martin Schutten. 2010. “The Effect of Primer-Template Mismatches on the Detection and Quantification of Nucleic Acids Using the 5’ Nuclease Assay.” The Journal of Molecular Diagnostics: JMD 12 (1): 109–17.

The Anopheles gambiae 1000 Genomes Consortium. 2020. “Genome Variation and Population Structure among 1142 Mosquitoes of the African Malaria Vector Species Anopheles Gambiae and Anopheles Coluzzii,” 1–14.

Untergasser, Andreas, Ioana Cutcutache, Triinu Koressaar, Jian Ye, Brant C. Faircloth, Maido Remm, and Steven G. Rozen. 2012. “Primer3--New Capabilities and Interfaces.” Nucleic Acids Research 40 (15): e115.

Wu, Jer-Horng, Pei-Ying Hong, and Wen-Tso Liu. 2009. “Quantitative Effects of Position and Type of Single Mismatch on Single Base Primer Extension.” Journal of Microbiological Methods 77 (3): 267–75.

Zajícková, K., A. Krepelová, and I. Zofková. 2003. “A Single Nucleotide Polymorphism under the Reverse Primer Binding Site May Lead to BsmI Mis-Genotyping in the Vitamin D Receptor Gene.” Journal of Bone and Mineral Research: The Official Journal of the American Society for Bone and Mineral Research 18 (10): 1754–57.

